# Sequence features of transcriptional activation domains are consistent with the surfactant mechanism of gene activation

**DOI:** 10.1101/2023.06.18.545482

**Authors:** Bradley K. Broyles, Tamara Y. Erkina, Theodore P. Maris, Andrew T. Gutierrez, Daniel A. Coil, Thomas M. Wagner, Caleb A. Class, Alexandre M. Erkine

## Abstract

Transcriptional activation domains (ADs) of gene activators remain enigmatic for decades as they are short, extremely variable in sequence, structurally disordered, and interact fuzzily to a spectrum of targets. We showed that the single required characteristic of the most common acidic ADs is an amphiphilic aromatic–acidic surfactant-like property which is the key for the local gene-promoter chromatin phase transition and the formation of “transcription factory” condensates. We demonstrate that the presence of tryptophan and aspartic acid residues in the AD sequence is sufficient for in vivo functionality, even when present only as a single pair of residues within a 20-amino-acid sequence containing only 18 additional glycine residues. We demonstrate that breaking the amphipathic α-helix in AD by prolines increases AD functionality. The proposed mechanism is paradigm-shifting for gene activation area and generally for biochemistry as it relies on near-stochastic allosteric interactions critical for the key biological function.

## Introduction

The expression of genetic information written in nucleotide sequences of DNA is the fundamental function of all living entities. The process is initiated by the binding of gene activators to corresponding gene promoters. In eukaryotes, gene activator proteins typically contain two obligatory parts: the DNA-binding domain (DBD), which is responsible for recognition and specific binding to the cognate promoter DNA sequence, and the activation domain (AD), which is responsible for the initiation of gene transcription. The two domains are drastically different in nature and in function. The DBD for each activator contains a certain conserved amino acid sequence and structure and binds to a specific consensus DNA sequence (Hossain et al., 2015; Luscombe et al., 2000). Contrarily, ADs are typically short, ranging from six to several dozen amino acids, and extremely variable in sequence, and it has been estimated that up to 10^24^ sequences are able to substitute for each other within the context of the same activator molecule (Abedi et al., 2001; Broyles et al., 2021; Erijman et al., 2020; Ma and Ptashne, 1987; Ravarani et al., 2018). In addition, ADs are intrinsically disordered in structure and bind to a wide variety of proteins and binding regions on the surface of these targets via “fuzzy”, low-specificity interactions (Mapp and Ansari, 2007; Sanborn et al., 2021; Tuttle et al., 2021). The mechanism of the function of ADs is a long-standing enigma, as their features defy the traditional specificity of sequence-to-structure-to-function paradigm of molecular biology.

Several models have been proposed to explain the mechanism of AD function. First is the traditional and widely accepted model of direct physical recruitment by ADs of coactivators and components of enzymatic transcriptional machinery. This model implies a certain level of specificity for AD sequences, such as a consensus sequence, or at least Short Linear Motifs (SLiMs); specific structural features, such as an amphipathic α-helix; and specific targets, for instance, the Mediator complex. However, recent high-throughput experimental data (Broyles et al., 2021; Erijman et al., 2020) and the results of bioinformatics analysis increasingly contradict the direct recruitment model, showing the lack of specificity of ADs at the sequence, structure, and target interaction levels. To alleviate this problem the recent development of the Liquid-Liquid phase separation (LLPS) concept suggests that during the transcriptional condensate formation the interaction partners - ADs, coactivators, and components of transcriptional machinery - are concentrated in coacervates, so that even low-affinity fuzzy interactions become possible (Boija et al., 2018). The role of ADs, as intrinsically disordered regions (IDRs) in this scenario is viewed often through lens of the “stickers and spacers” idea (Choi et al., 2020; Staller et al., 2022), whereby the hydrophobic residues (stickers) of IDRs are kept in solvent exposed active configuration by hydrophilic, often acidic, residues (spacers), thus serving as seeds for 2D prewetting (Renger et al., 2022) and 3D condensation processes (Boija et al., 2018). Within the LLPS models function of ADs is still traditionally considered within the frame of the AD-coactivator interactions, which is essentially an extension of the recruitment model (Boija et al., 2018; Staller et al., 2018; Staller et al., 2022).

Another, newer and not yet widely accepted model (Broyles et al., 2021; Erkine, 2018) considers acidic ADs as amphiphilic surfactants with aromatic and acidic extremities that act on the interface between DNA and histones of promoter nucleosomes. Presumably, by intercalation of aromatic moiety between bases of DNA and by acidic residues interfering with salt bridges between DNA phosphates and histone tails, ADs trigger chromatin remodeling leading to the formation of nucleosome-free regions, which in turn, stimulate formation of transcription condensates. The surfactant model postulates the extremely high variability of AD sequences, because the requirement for the functional sequence is the presence of only a limited number of acidic and aromatic amino acids. The model requires no specific structure but spatial and structural flexibility and presumes near-stochastic interactions with targets at the anchoring site (i.e., the gene promoter). The absence of a specific sequence or structure, and a near-stochastic target interaction mechanism, of a protein domain that is critical for a fundamental biological function presents a new paradigm, challenging the traditional foundational biochemical principle of specificity rooted in the sequence-to-structure-to-functional-interaction triad.

Here, we provide experimental data that allow us to compare and possibly discriminate between proposed mechanistic models. By synthesizing and testing large sets of specific AD sequences *in vivo*, we demonstrate that, in line with the surfactant model and contrary to the specificity required by the recruitment model, the presence of only one tryptophan and aspartic acid, each represented as a single side chain-bearing amino acid residue within the AD region, is sufficient for gene activation. Also contrary to the expectations of the recruitment model, we demonstrate that the presence of the amphipathic α-helix structure is detrimental for AD functionality, while breaking this structure by insertion of prolines significantly increases AD functionality, providing a rationale for the existence of the entire class of proline-rich ADs. With these and other data, and in line with the surfactant model, we hope to change the gene expression paradigm and establish near-stochastic interactions as required for, rather than detrimental to, this and possibly other crucial biological functions.

## Results

To elucidate the mechanism of AD function, we developed a high-throughput assay to test the *in vivo* functionality of individual sequences within a library of 12,400 synthetically generated sequences. These sequences were designed to address multiple questions regarding potential determinants of AD function, such as length, composition, structure preference and others. The synthesized library was then cloned into a yeast centromeric shuttle vector, fusing each AD sequence to the Gal4 DBD. Then, the library was transformed into the yeast two-hybrid Y2HGold tester strain, and the transformed yeast were screened for growth, which was dependent on the expression of the *Gal*-Aureobasidin antibiotic resistance reporter gene (Fig. 1A). The library DNA for different growth time points was isolated and sequenced to determine the number of reads for each individual sequence and its change over time. The growth slope for each sequence was calculated and served as a measure of the AD functionality. Since the results were obtained within the scope of the whole library pool screening, the results for individual sequences and for all different sequence sets can be considered to be obtained under identical experimental conditions and thus to be accurately comparable. In addition, each sequence was labeled by multiple individual barcodes; thus, the results for each individual sequence are the mean of multiple (typically five) independent experimental repeats. As part of the library we used the sequences of known AD regions, such as Gal4(840-857), Gal4(860-872) and VP16 sequences (Piskacek et al., 2007; Wu et al., 1996), as internal positive controls (Fig. 1B). As negative controls, we used a null sequence that contained a stop codon after the DBD and a sequence containing a stretch of 20 glycines (G), which was shown to be neutral for AD functionality under similar experimental conditions (Broyles et al., 2021). To ensure high stringency, the cutoff for a “functional AD” was defined as the mean of the five highest values produced by 50 independent stop codon-null sequences.

**Figure 1.**
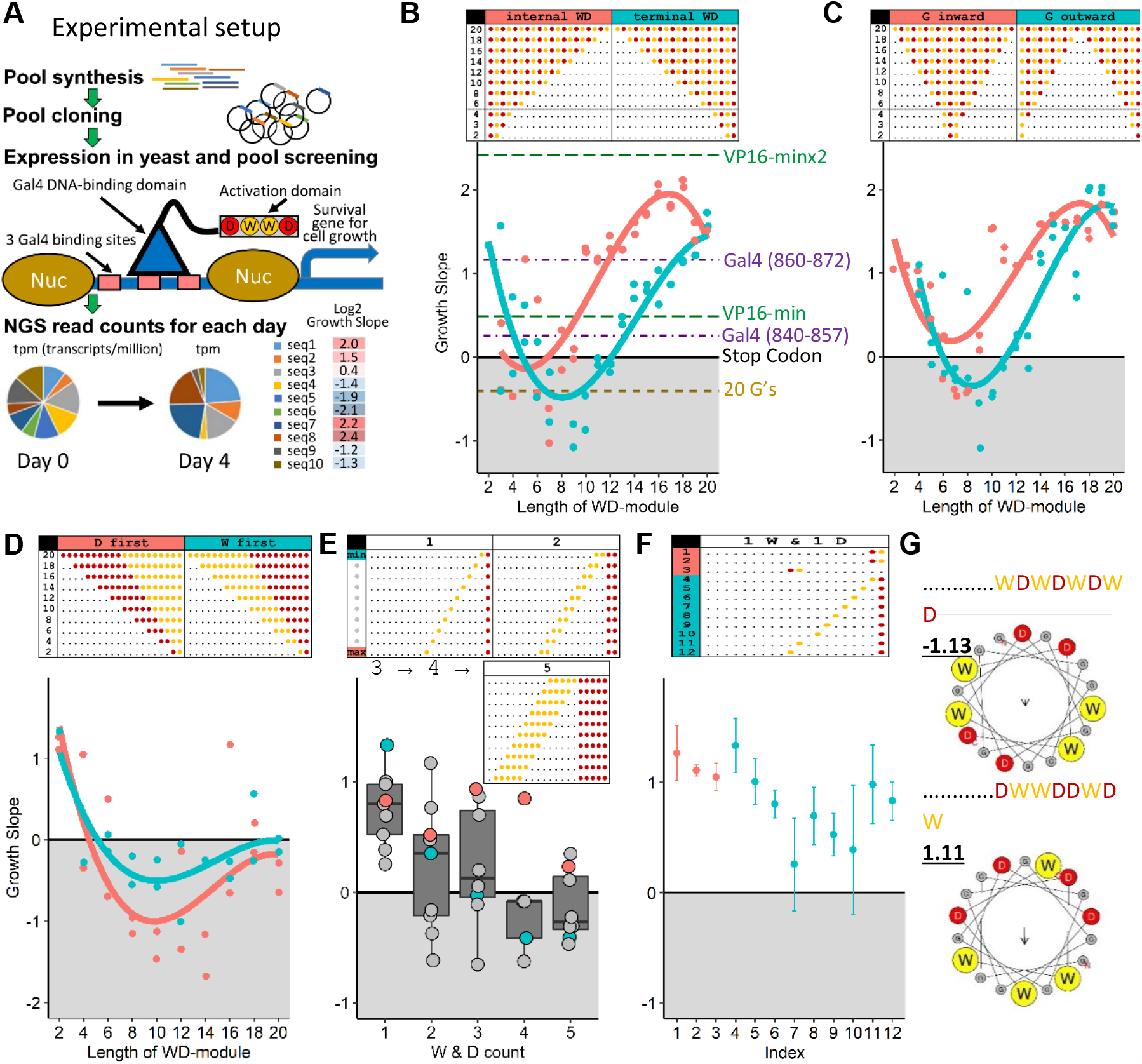
A single W and a single D are sufficient for the functionality of AD. **A** – Experimental setup: oligo pool synthesis, followed by cloning in bacteria, then isolation of plasmid library and transformation in yeast, followed by screening for growth phenotype determined by expression of the reporter gene regulated by the activator with a specific AD, then isolation of DNA pool, NGS sequencing, and data analysis (for more details see Methods section and Supplementary Figure 1). **B** – Growth of sequences with different numbers of WD repeats. X axis: individual sequences indicated in the inset table, where black dots represent glycine, yellow dots represent tryptophan, and red dots represent aspartic acid residues. Y axis: Log2 growth slope. Axes are same in C, D, E, F. **C** – Sequences with different numbers of WD repeats, either surrounded or interrupted by repeated glycines. **D** – Sequences with non-alternating clusters of Ws and Ds. **E** – Sequences with non-alternating clusters of Ws and Ds, separated by varying numbers of Gs. **F** – Sequences with a single W and D, separated by varying numbers of Gs. Error bars show growth slope +/- root-mean-square-deviation (RMSD) of the fit of the growth slope. **G** – Log2 growth slopes and images of the α-helix frontal view for two different sequences with four W’s and four D’s, showing that despite identical composition, only one is functional.

### A single aspartic acid and single tryptophan residue within the AD region are sufficient for functionality

Previous reports indicated that the most beneficial residues for AD functionality are aromatic and acidic amino acids, and the highest gain in prediction of AD functionality using machine learning on large >10^6^ set of diverse random sequences was derived from the presence of amino acids W and D (Broyles et al., 2021). In addition, high AD functionality was repeatedly demonstrated for the monotonous WDWDWDWDWDWDWDWDWDWD sequence(Ravarani et al., 2018), indicating that for high functionality, it is sufficient to have only W and D. Here, we analyzed sequences with different numbers of WD repeats to determine how functionality depends on this feature and found that decreases in WD repeat number generally correlate with decreases in functionality (Fig. 1B). Unexpectedly, the sequence GGGGGGGGGGGGGGGGGGDW, containing just one W and one D, showed residual functionality (1.11±0.02 [95% CI]) comparable to that of the VP16 minimal AD module (0.55±0.28). Similarly, analysis of WD sequences surrounded by G residues or WD blocks separated by a stretch of G residues (Fig. 1C) revealed functionality for sequences containing a single W and a single D. The lowest functionality was demonstrated for sequences containing four or five Ws and Ds, and the highest functionality was demonstrated for the DWx9 sequence (2.12±0.22). The answer to the same question about the amount of Ws and Ds sufficient for AD functionality within the context of sequences with non-alternating clustered Ws and Ds (Fig. 1D) is that one W and one D is sufficient for at least a low level of functionality. Sequences with a single W and single D separated by different numbers of Gs (e.g., WGD, WGGD, WGGGD, see Fig. 1E and F) are also functional. By contrast, sequences containing more than two adjacent Ws or Ds (e.g., GGGGGGGGGGGGGGGGWWWDDD or GGGGGGGGGGGGGGGGDDDWWW) are nonfunctional, and the sequence functionality generally drops with increasing lengths of homo-W and homo-D stretches (Fig. 1D). These results are in good agreement with previous observations that clustering of separate acidic and separate aromatic residues within the AD region is detrimental to functionality (Broyles et al., 2021). Interestingly, some functionality was observed for sequences containing both W homostretches and D homostretches separated from each other by larger numbers of Gs (Fig. 1E, red dots). In examining why the sequence containing the WDWDWDWD block is nonfunctional, we found that DWWDDWDW, a different sequence with an identical composition, is functional (Fig. 1G). A possible explanation for this observation is that while and if forming a 3D α-helix, Ws within the WDWDWDWD sequence are forming pi-pi interacting pairs, while in DWWDDWDW, at least one W remains free. That in turn suggests that solvent exposure of at least one aromatic residue in the AD sequence is required for functionality and is consistent with the finding that a single D and single W is sufficient for functionality (Fig. 1F).

### Balance and intermixing of acidic and aromatic residues underlie AD function

To expand the analysis, we turned to the other part of our library, which contains a set of sequences with all possible combinations of W and D at 12 positions (3968 quantified out of 4096 possible sequences). The *in vivo* screening revealed 1330 functional and 2638 nonfunctional sequences within this set. Analyzing this set as a whole, we found that in general, two features are important for functionality: the balance between W to D residues in the sequence, and how they are intermixed (Figs. 2A and 2B). Similar patterns were noticed when the balance and intermixing between aromatic and acidic residues were analyzed within a much larger set of ADs in the context of the Gcn4 activator or HSF activator (see reference (Broyles et al., 2021) and Fig. 2C).

**Figure 2.**
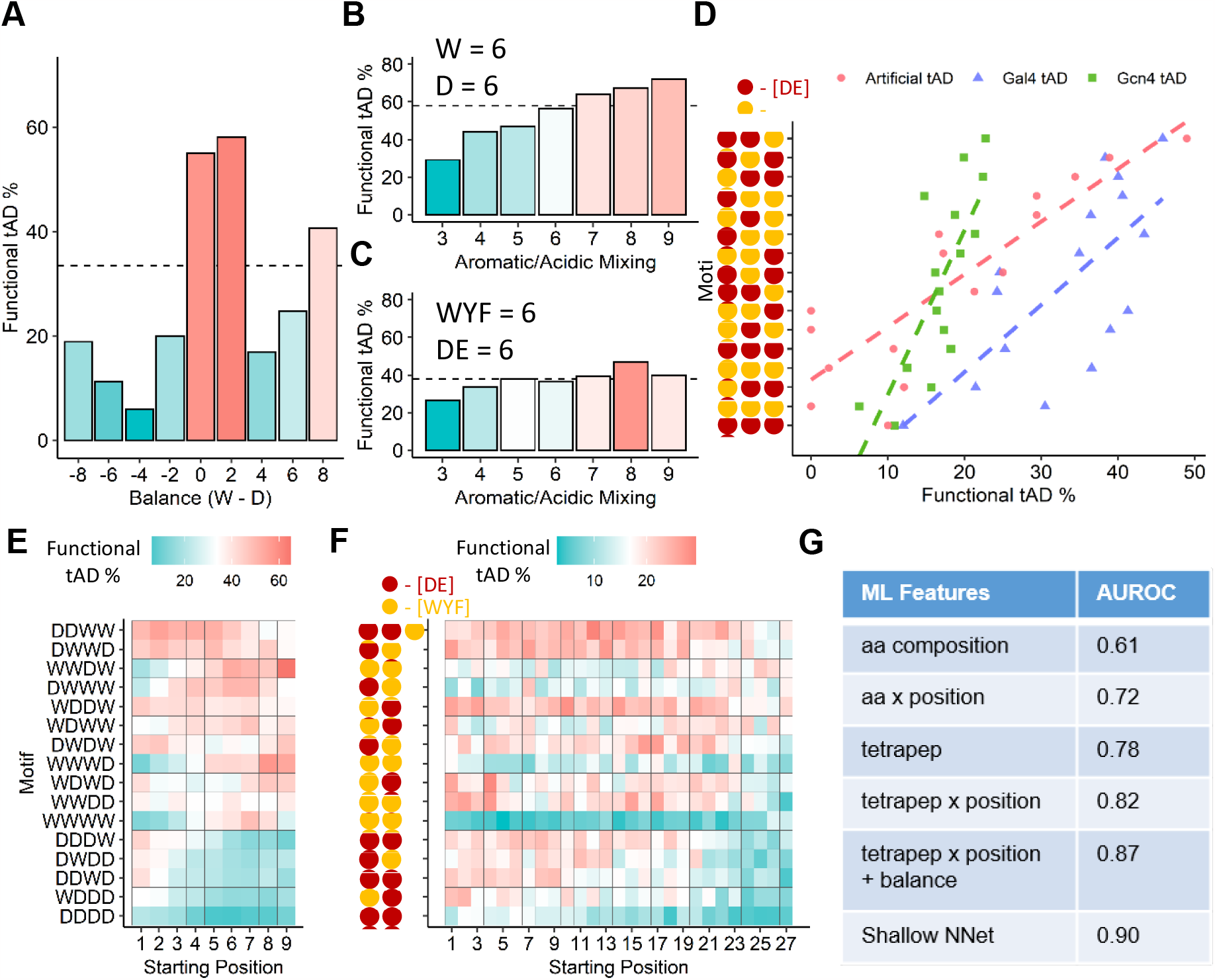
The balance and intermixing of acidic and aromatic residues is beneficial for AD function. **A** – X axis: balance score, calculated as Balance=n(W)-n(D); Y axis: % of functional sequences in the sub-library of sequences containing all combinations of W and D for 12 positions (WD12 library, 3968 sequences quantified). The horizontal dashed line represents the mean functional AD percent for the library. **B** – X axis: mixing score, calculated as Mixing=n(WD)+n(DW); Y axis: % of functional sequences in the set of sequences containing all combinations of 6 W and 6 D (906 sequences quantified). **C** – X axis: mixing score, calculated as Mixing=n([WYF][DE]) + n([DE][WYF]) for the previously published AD dataset screened within the Gcn4 context ^6^; Y axis: % of functional sequences in the set of sequences from the Gcn4 random peptide library with 6 [WYF] and 6 [DE] (3018 sequences total). **D** – X axis: % functionality of sequences that contain the specified tetrapeptide motif; Y axis: tetrapeptide motifs. Regression lines are provided to demonstrate concordance between the three libraries, and motifs were ordered based on average % functionality between the three libraries, with the most functional on top. **E** – X axis: Starting amino acid position of tetrapeptide in tAD module for the WD12 library; Y axis: 16 sequence combinations for tetrapeptides containing D and W, Tile fill: % functionality of sequences that contain the specified tetrapeptide motif at the indicated position. Motifs were ordered by overall % functionality. **F** – X axis: Starting amino acid position of the tetrapeptide in the tAD module for the Gcn4 library ^6^; Y axis: 16 sequence combinations for tetrapeptides containing [DE] and [WYF], Tile fill: % functionality of sequences that contain the specified tetrapeptide motif at the indicated position. Motifs order is the same as in panel E. **G** – ML accuracy on the reserved testing set (20% of WD12 library) and trained on 80% of WD12 library, measured as area under the receiver operating characteristic (AUROC).

To examine whether these functional sequences might contain a specific mini-motif, we tested all 16 WD tetrapeptide variants and found that tetrapeptide sequences containing an excess of Ds, especially Ds clustered together, are generally detrimental, while D and W intermixing is beneficial and creates a number of functional tetrapeptide sequences, with DDWW as the top performing sequence (Fig. 2D). A very similar trend of aromatic-acidic tetrapeptide motif distribution is obvious from our analysis of two independent *in vivo-*tested AD and previously published sequence datasets created on the basis of natural AD sequences in the context of artificial DBD-ER fusions (Sanborn et al., 2021) and a sampling of large unbiased random sequence ADs in the context of Gcn4 (Erijman et al., 2020). This similarity of the trends suggests an activator-independent general mechanism for AD function.

To test whether the position of the tetrapeptides within the AD sequence is important, we calculated the probability of each tetrapeptide contributing to functionality when positioned in different parts of the AD region (Fig. 2E). This analysis indicated that while W and D intermixing is beneficial, W-rich sequences are generally more beneficial at the spatially freer end of the molecule, while D-rich sequences are beneficial internally. Similar trends are observed for the broader set of acidic-aromatic tetrapeptides in the Gcn4 context (Fig. 2F and reference (Broyles et al., 2021)). When we used individual tetrapeptides, their position within the sequence, and the balance between acidic and aromatic residues as features for regression ML model training, we observed that each feature had a positive value for the prediction of AD functionality, and the combination of all these features produces the most accurate prediction (Fig. 2G).

The position effect is much clearer when the functionality contributions of W and D within the AD sequence space are analyzed directly (Fig. 3A). Generally, the positive contribution of W increases when it is positioned toward the end of the molecule, while D shows the exactly opposite behavior. This observation is consistent with our previous analysis (Broyles et al., 2021); however, the trend breaks at the last two positions within a 12-amino-acid AD. In examining what sequences display higher functionality with generally detrimental terminal D(s), we found that it is especially beneficial if a cluster of Ws precedes the D(s) (Fig. 3B). A similar trend was observed when we analyzed the acidic and aromatic amino acid distributions within a previously published (Erijman et al., 2020) dataset of random AD sequences in the Gcn4 context (Fig. 3C). Comparing the different sequences containing a cluster of 5 Ws, which is usually detrimental for functionality, we found that flanking such clusters with Ds is generally beneficial, with the highest functionality observed if the majority of Ds are situated internally (Fig. 3D). Molecular modeling suggested that the D-flanking effect likely occurs because the repelling charges of aspartic acid residues prevent the tryptophan moieties from forming a hydrophobic mini-globule or a disordered aggregate supported by hydrophobic and pi-pi interactions between aromatic rings (Fig. 3E).

**Figure 3.**
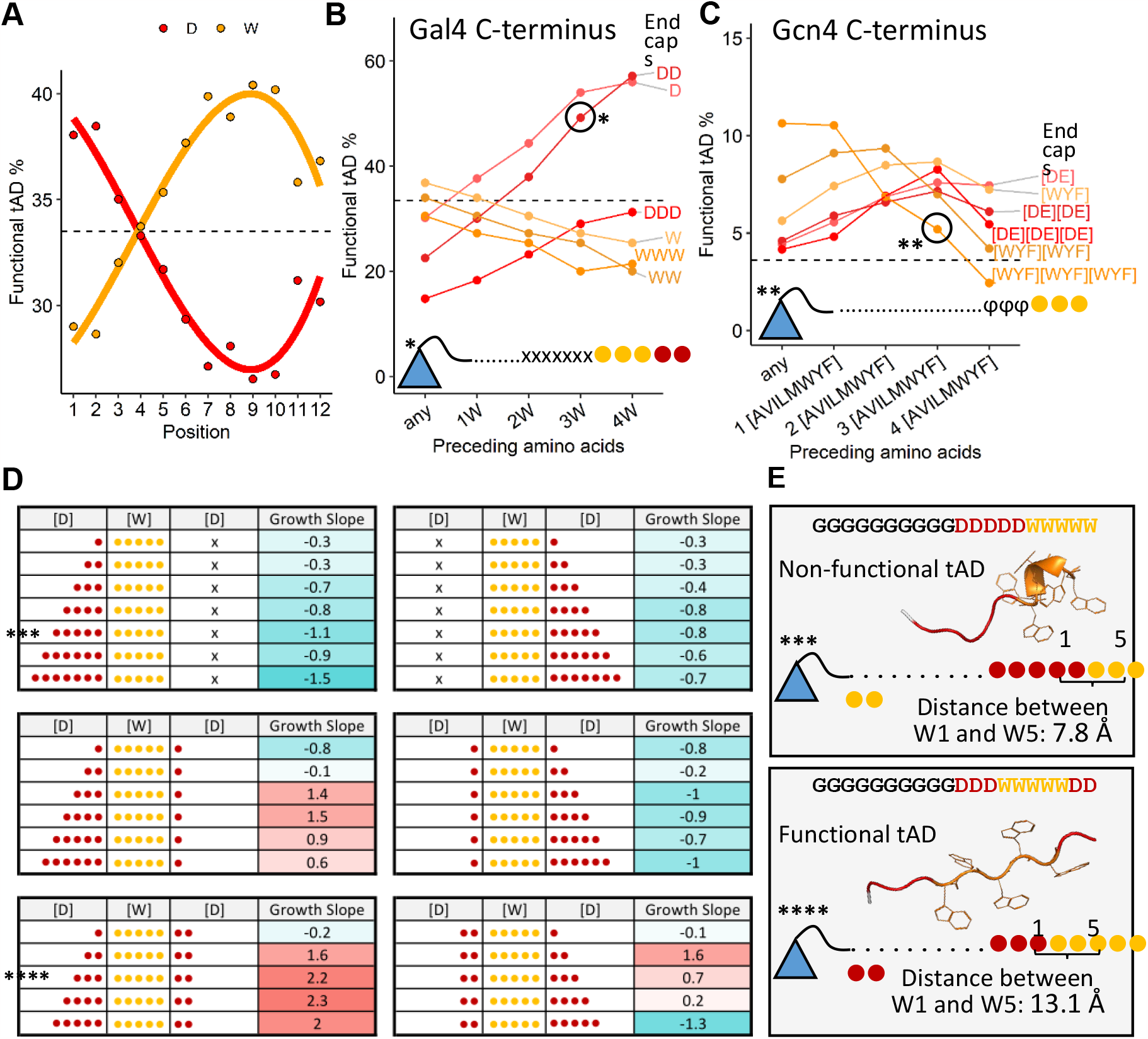
Ws are generally beneficial at the end of the molecule, while Ds – internally, with the exception when Ds rescue the AD functionality by flanking adjacent Ws in the sequence. **A** – X axis: position of D (red) or W (yellow) within the sequence; Y axis: % functionality of sequences in the sub-library of sequences containing all combinations of W and D for 12 positions (3968 sequences quantified). **B** –X axis: size of W cluster preceding indicated end cap for sequences representing each line; Y axis: same as in A. (*) sequence construct shows tAD constructs of indicated sequence where “.” = G, “x” = [DW], 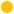 = W, and 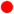 = D. **C** – same as in B, calculated for the Gcn4 library ^6^ using [WYF] instead of just W and [DE] instead of just D for end caps. (**) sequence construct shows tAD constructs of indicated sequence where “.” = any AA, ϕ = [AVILMWYF], and 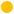 = [WYF]. **D** – Growth slopes of sequences with different numbers of Ds flanking a stretch of 5 Ws. 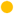 = W, and 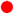 = D. **E** – sequence constructs of (***) and (****) sequences from panel D, “.” = G, 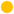 = W, and 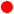 = D. AlphaFold2 predicted structures shown for tAD region. Distance between α-carbon of first tryptophan (W1) and last tryptophan (W5) was measured from predicted structure.

### Formation of an amphipathic α-helix may be detrimental to AD functionality, while breaking the helix with proline increases the gene activation potential

Since folding of the AD sequence into a specific structure (Fig. 1-3) seems to play an important role, and because the amphipathic α-helix is specifically considered as an important structural feature of ADs (Erijman et al., 2020; Sanborn et al., 2021; Staller et al., 2018; Tuttle et al., 2021), we created a library of sequences all containing 5 Ws and 5 Ds interspersed with random amino acids (WxDxWxDxWxDxWxDxWxD, henceforth called the WD5 library). Any sequence of the WD5 library, if folded into the canonical helix structure, creates an amphipathic α-helix (Fig. 4A). Even though the probability of the α-helix formation in this context is negatively affected by the repulsion of Ds on one side of the α-helix, the specific amino acids in the X positions potentially can facilitate or exacerbate the α-helix formation in individual sequences. WD5 library in vivo screening followed by DNA sequencing, normalization of the number of reads for each sequence to that at the 0-time point, and AD functionality cutoff based on redundant stop codon null sequences, as described above, confirmed that of 107,975 distinct sequences tested, 19.6% were functional ADs. We binned the sequences by their predicted percent α-helix formation and found that the percentage of functional sequences in each bin decreased as the prediction of the α-helix fraction increased (Fig. 4B). By analyzing sets of sequences enriched with individual amino acids within the WD5 library (Fig. 4C), we found that increasing the number of basic amino acids between Ws and Ds was detrimental, which was consistent with the highly negative effect of K and R on AD functionality (Broyles et al., 2021). A similar negative effect was observed for sequences enriched with additional (> 5) aromatic residues and to a lesser degree among sequences enriched with additional acidic residues, which is consistent with the results of Fig. 2A and the previously observed negative effect of shifting the balance between acidic and aromatic residues (Broyles et al., 2021). Unexpectedly, a progressive increase in the number of proline residues (P) within the WD5 sequences was correlated with a significant increase in the probability of functionality, from 15.7% with zero prolines to 55.6% with five prolines (Fig. 3C). Proline residues are known to be potent helix breakers. Thus, breaking the amphipathic α-helix of the WD5 sequence is generally beneficial for functionality of the sequence within the WD5 context. This proline effect was not observed for sequences from random-sequence libraries (Erijman et al., 2020; Ravarani et al., 2018), suggesting that the positive proline effect may be specific to the amphipathic helix context of the WD5 library. The presence of prolines in this case potentially prevents tryptophan rings from interacting with each other on one side of the amphipathic helix, thus keeping the Ws exposed to the solvent (Fig. 4D). Fig. 4E confirms the phenotypes of representative sequences including a proline-rich sequence (indicated in red) discussed above.

**Figure 4.**
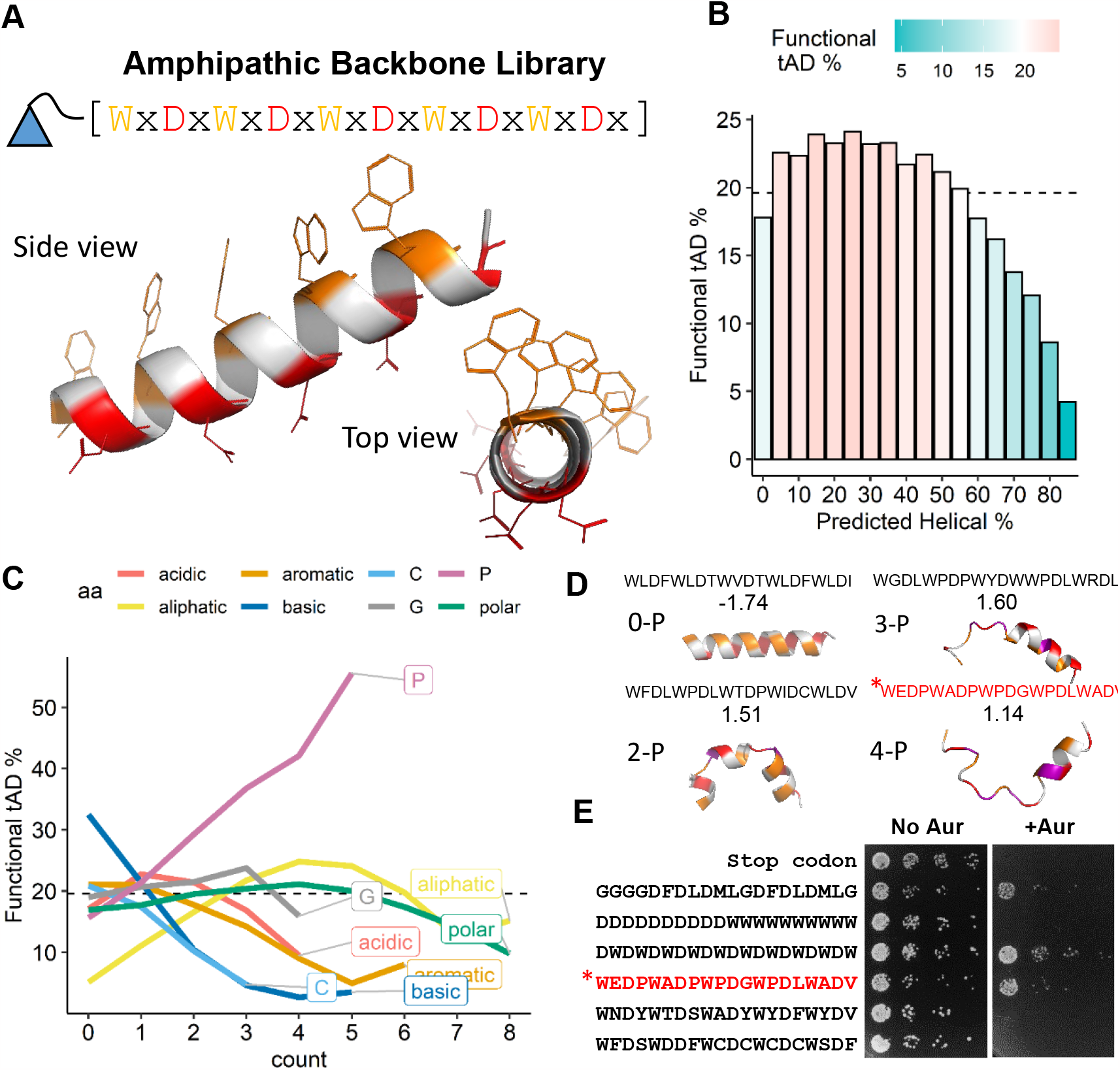
Formation of the amphipathic α-helix is generally detrimental for AD functionality, and insertion of proline is beneficial. **A** – graphical representation of sequences for the WD5 library (107975 sequences quantified): each member of which has five Ws, five Ds, and ten random amino acids represented by ‘x’ between Ws and Ds. **B** – X axis: % α-helix predicted by the SPOT-1D algorithm for each set of sequences from the WD5 library; Y axis: % functionality of the set of corresponding sequences in the WD5 library. **C** – X axis: count of corresponding amino acid residues between set Ws and Ds of WD5 library; Y axis: % functionality of the set of corresponding sequences. Amino acid groups: acidic [DE], aliphatic [AVILM], aromatic [WYF], basic [RHK], special [CGP] not grouped, polar [STNQ]. **D** – Growth slopes with 3D structures of sequences with varying numbers of proline residues predicted by AlphaFold2. **E** – Growth phenotype on media with and without aureobasidin for cells expressing the indicated representative sequences. Spots are conglomerates of yeast colonies representing threefold serial dilutions of corresponding cell cultures.

## Discussion

The enigma of the AD mechanism is recognized for decades (Strader et al., 2023), and it has been studied extensively since the invention of high throughput experimentation (Erijman et al., 2020; Erkine, 2018; Ferrie et al., 2022; Sanborn et al., 2021). The main challenge seems to be that ADs do not follow the conventional sequence-to-structure-to-interacting partners paradigm and engage in numerous, often uncertain “fuzzy” interactions, either within the target (Tuttle et al., 2018) or between multiple different targets (Aguilar et al., 2014; Ansari et al., 2002; Bhaumik et al., 2004; Brown et al., 2001; Capella et al., 2014; Chabot et al., 2014; Hermann et al., 2001; Jeong et al., 2001; Keung et al., 2014; Khan et al., 2011; Knutson and Hahn, 2011; Neely et al., 2002; Neely et al., 1999; Ozer et al., 1996; Prochasson et al., 2003; Qiu et al., 2004; Stargell and Struhl, 1995; Tan et al., 2000; Tuttle et al., 2018; Warfield et al., 2014). Considering the extreme variability of the AD sequences (up to ∼10^24^ variants for 20 amino acid size and 1-3% of functional ADs for random library screens (Erijman et al., 2020; Ravarani et al., 2018)), fuzzy interactions with targets are likely of extremely low specificity and affinity. That creates difficulty in applying the model of the coactivator recruitment mechanism for ADs. Recognizing this problem in the recent development of the LLPS concepts, new ideas were proposed to solve the long-standing enigma. The formation of super-enhancer condensates brings the AD and potential interacting targets in proximity and increases local concentrations of interacting partners allowing otherwise unlikely interactions to happen (Boija et al., 2018). The formation of coacervates recently was suggested to be facilitated and regulated by the prewetting processes (Morin et al., 2022) whereby the local increase of charged residues and exposure of individual hydrophobic amino acids creates 2D surfaces conducive to the 3D condensates formation. However, the biological stimuli of prewetting and coacervate formation/dissociation remains obscure and the coactivator recruitment mechanism for ADs remains a dominant conventional mechanism even within the LLPS paradigm.

The competition of multiple 2D prewetting and 3D LLPS processes is dynamic. From the transcription activation perspective, the main competing phases are dissociation/formation of hetero/euchromatin (Larson and Narlikar, 2018; Zhang et al., 2022) and formation/dissociation of transcriptional machinery condensates (Boija et al., 2018). The AD sequences of our study are intimately involved in all these activities. The properties of AD sequences we identified mostly fall into and described by the “stickers and spacers” concept whereby hydrophilic “spacers” expose hydrophobic “stickers” critical for the multivalent interactions conducive to LLPS (Choi et al., 2020). However, the AD sequences in our results, and many other ADs, are very short (Fig. 1) to the extent that positive AD activity can be observed even when the AD region contains all sidechain-less glycine amino acid residues and only a single aspartic acid and a single tryptophan residue (Fig. 1C). In this case the amount of “stickers and spacers” is minimal, likely limiting the effect of the AD action to the trigger rather than to action as a major organizer of 2D prewetting seeds or full-size 3D condensates.

The minimalistic ADs are highly compatible with the acidic/aromatic surfactant model of the AD action (Broyles et al., 2021; Erkina and Erkine, 2016; Erkine, 2018; Ravarani et al., 2018). According to this model, ADs trigger local chromatin remodeling events driven in bulk by ATP-dependent enzymatic chromatin remodels and histone chaperones, thus leading to the exposure of multiple charged phosphate groups of local naked DNA gene promoter regions. The AD-triggered initiation of local chromatin phase transition and exposure of the multivalent charged DNA backbone in turn can stimulate 2D prewetting and 3D transcription condensate formation (Hnisz et al., 2017; Morin et al., 2022; Wei et al., 2020). In this model ADs, instead of serving as coactivator recruitment entities, which is highly unlikely due to only having single acidic and single aromatic residues present (Fig. 1F), likely serve as triggering surfactants (Broyles et al., 2021; Erkine, 2018). It is also worth noting that in our study, although we analyzed designed AD sequences created by the cutting-edge massive parallel synthesis method, and thus the sequences could be characterized as “synthetic” or “artificial”, all of the sequences were verified in vivo and thus represent bona fide ADs. In addition, naturally occurring AD sequences such as VP16 and Gal4 AD modules were included in the library as internal controls.

Consistent with both the “stickers and spacers” (Choi et al., 2020) and the surfactant model (Broyles et al., 2021; Erkine, 2018) is our observation that the introduction of increasing number of prolines in a 20 amino acid stretch of AD region leads to a significant increase of gene activation (Fig. 4C). The WD5 library sequences all, if folded into an α-helix, should form an amphipathic structure, which is a structural element that is traditionally considered as important in the recruitment of coactivators and transcriptional machinery by the AD (Erijman et al., 2020; Sanborn et al., 2021; Staller et al., 2018; Tuttle et al., 2021). The amphipathic α-helix in this case fits the valley or even a tunnel of the AD binding site on the coactivator surface, thus ensuring multiple bonds necessary for the recruitment event (Sanborn et al., 2021). However, the results of our analysis of thousands of sequences suggest that the presence of the amphipathic α-helix in a WD5 AD is, if not detrimental, then at least not beneficial for function, and breaking this structure by prolines increases the probability of AD functionality proportionally to the increase in proline content (Fig. 4C). Consistent with the “stickers and spacers” and surfactant models, breaking the structure and thus making Ws and Ds more solvent-exposed and more available for interaction with target(s) is functionally beneficial. Following the same logic, adding a surplus of Ws in a WD5 sequence increases the likelihood of pi-pi interactions between neighboring Ws, thus likely leading to the formation of a locked noncanonical structure. This interpretation is also consistent with the observation that flanking W clusters with Ds restores the functionality of otherwise inactive sequences with multiple Ws, which, without breaking them with Ds, are likely to collapse into intermolecular pi-pi interaction stabilized aggregates (Fig. 3D and 3E). The gain of functionality for sequences with identical composition correlates with the ability to adopt a structure that ensures an individual tryptophan maintaining a solvent exposed configuration.

The positive effect of proline, demonstrated for 830 sequences with ≥3 prolines within the WD5 amphipathic α-helix context, suggests the explanation for existence of the entire proline-rich class of ADs. This class was described several decades ago (Mapp and Ansari, 2007; Mitchell and Tjian, 1989), but the reason for the functional preference of proline in ADs has remained obscure. Considering the “stickers and spacers” and the surfactant models, proline residues likely prevent internal aggregation of Ws and ensure the exposure of Ws and Ds, for interactions with targets. Following the same logic, 32% of sequences with multiple representation of Ws and Ds are functional (2579 functional AD sequences out of 8094 sequences with ≥2W and ≥2D in our design library), likely because of internal repulsion of Ds, which disrupts conventional structures or non-conventional aggregation of Ws, maintaining aromatic residues in the solvent exposed thus functional configuration.

Another important observation is the general preference for aromatic residues at the spatially free terminus of the molecule observed in this study (Fig. 3A) and previously (Broyles et al., 2021; Erijman et al., 2020). While for the recruitment model, initial interactions with the target are based on scanning by the negatively charged acidic residue and establishing a strong initial salt bridge with the target (Ferreira et al., 2005), for the surfactant model the negative charge at the end is disadvantageous for function due to the repulsion of DNA phosphates and hence the interference with the required initial intercalations of aromatic residues into DNA. Contrary to the recruitment model (Ferreira et al., 2005) and consistent with the surfactant model, we demonstrate that the exception from the aromatic end preference is observed only when a terminal acidic residue(s) is(are) required for the unraveling of aromatic clusters (Fig. 3).

Although the role of acidic residues in exposing aromatics is important from the perspective of “stickers and spacers” model, the main function of acidic moieties in the surfactant model is the proposed interference in DNA-histone nucleosome salt bridges. This action of the amphiphilic AD triggers promoter chromatin remodeling, freeing the promoter DNA (Erkine, 2018). The attraction of multiple transcription machinery components to the exposed promoter DNA may explain the liquid–liquid phase separation (LLPS) observed in the eukaryotic nucleus upon induction of transcription (Boija et al., 2018).

Even though we emphasized a possibility of ADs functioning as triggering surfactants, once the phase separation occurs, the AD can assume a more traditional role within the condensate interacting with the transcriptional machinery components by fuzzy interactions stabilizing, for instance, Mediator complex (Sanborn et al., 2021; Tuttle et al., 2018; Tuttle et al., 2021) and other components within the condensate (Chen and Pugh, 2021). Another traditional AD function with the condensate that requires further investigation, especially in higher eukaryotes, can be a role of ADs in release of posed polymerase (Henninger et al., 2021).

Although suggesting the solution for the decade’s long enigma of ADs and potentially filling the long-standing knowledge gap in our understanding of gene expression mechanisms, the investigation of the surfactant model is still in its initial phase. The limitation of our study is that the data supporting the surfactant model is the result of the in vivo experimentation, which requires a systematic in vitro follow up study examining the possibility of the surfactant action of ADs on the DNA-histone interface of the nucleosome, and consideration of those results within the frame of both recruitment and surfactant models. Considering the extremely transient nature and low affinity/specificity of AD interactions illustrated by this study and others (Broyles et al., 2021; Erijman et al., 2020; Sanborn et al., 2021) designing these in vitro experiments is not trivial, as it requires new assays based on the acceptance of near-stochastic surfactant-like interactions as functionally relevant. These systematic in vitro studies are outside of the scope of the current study and likely will be the focus of future investigations.

While the surfactant model allows us to look at the most important function in biology – gene expression – from a perspective of low affinity/specificity interactions of IDRs, it is not the only biological function with an unexplained mechanism that requires close attention. Near-stochastic interactions and IDRs have been shown to play important roles in such processes as mRNA processing, apoptosis, molecular transport within and between cells, glycolysis, and many others (Bondos et al., 2021; Oldfield and Dunker, 2014). Breaking from the specific sequence-to-structure-to-function paradigm and considering near-stochastic interactions as fundamentally important and not detrimental opens an avenue to a completely new branch of biochemistry and molecular biology (Erkine, 2018).

## ACKNOWLEDGMENTS

We thank Marcos Oliveira for discussions and suggestions. The work was supported by NSF grants MCB 1925646 (to A.M.E).

## AUTHOR CONTRIBUTIONS

A.M.E conceived the project. T.Y. E. performed in vivo part of the work. B.K.B, A.T.G, T.P.M, D.A.C, T.M.W, and C.A.C performed data analysis. C.A.C oversaw methods and visualizations. A.M.E wrote the manuscript. All authors edited and approved the manuscript.

## DECLARATION OF INTERESTS

The authors declare no competing interests.

## STAR Methods

### Library construction, cloning, and screening

The parental library plasmid was constructed by cloning the fragment containing the *ADH1* promoter and the Gal4(1-147) DBD cassette, PCR amplified from the commercially available pGBKT7 vector. The PCR fragment was cloned into the *SacI* and *KpnI* sites of the centromeric yeast shuttle vector pRS314.

The design library containing 11,500 individual sequences, each with an individual 20-nucleotide barcode directly following the stop codon to improve alignment performance, was synthesized at the GenScript commercial facility, amplified by PCR five times (each time appending a unique BioRep barcode), quantitated for the DNA content, and mixed in equal proportions into a single pool. For description of more detailed steps, see supplementary Fig.1. The pool was cloned into the *NcoI* and *SalI* restriction sites remaining from the pGBKT7 fragment of the parental library plasmid. The library complexity was estimated by individual colony counts after transformation for a fraction of the total transformation mix, then multiplying by the fraction factor. Total complexity was estimated to be ∼10^6. The total content of individual sequences within the library was determined by NGS at GenScript. The NGS sequencing also confirmed the in-frame fusion of AD sequences to the Gal4 DBD region. After the bacterial cloning and verification, the plasmid library was isolated for the following yeast transformation.

At the Butler University research lab, the isolated plasmid library was transformed into the yeast strain Y2HGold, available commercially from Clontech/Takara. The maintenance of the library complexity was determined by the individual colony count for a fraction of a transformation mix as described above for the bacterial transformation. The number of individual yeast transformants for entire library was estimated to be ∼10^6. After transformation, the whole-library cell culture was transferred into the –trp synthetic yeast growth medium containing 200 μg/ml of aureobasidin and grown for four days with daily 1/100 dilution to maintain the culture in the mid-log phase. Cell culture samples were taken at 0, 1, 2, 3, and 4 days. DNA was isolated using a Thermo Scientific Pierce Yeast DNA Extraction Reagent Kit. The library component was isolated by PCR using the Invitrogen AccuPrime SuperMix I kit with primers containing Illumina adapters and barcodes unique for each DNA sample. DNA samples were controlled for purity, repeatedly quantitated for DNA content, and sequenced at the NovoGene commercial facility.

The semirandom WD5 library, containing sequences encoding peptides with five Ws and five Ds separated by random amino acids, was constructed from oligonucleotides synthesized at the IDT commercial facility according to the target sequence: ATCTCAGAGGAGGACCTGCATATGGGATGGNNNGATNNNTGGNNNGATNNNTGGNNNGATNNNTGGNNNGATNNNTGGNNNGATNNNTAGGTAGCTATGCGACCTGCAGCGGCCGCATAACTAGCATA where Ns are random nucleotides forming a triplet for a random amino acid.

At the GenScript commercial facility, the oligo was converted into the double-stranded form, digested with *NcoI* and *SalI*, and cloned into the corresponding restriction sites of the parental library vector, as described for the design library. The library complexity was estimated as described above by individual colony counts after transformation and assessed to be ∼10^6. The insertions and in-frame fusions with Gal4 DBD were confirmed by PCR and Sanger DNA sequencing for 40 randomly chosen individual plasmid isolations. After the bacterial cloning and verification, the entire plasmid library was isolated for the following yeast transformation.

The isolated WD5 library was transformed into yeast Y2HGold strain, the maintenance of the library complexity was confirmed by the individual colony count for the fraction of the transformation mix, as described above. Screening procedure, sample preparation, and NGS sequencing were also the same as for the design library.

### Sequence read processing

The reads from the semirandom WD5 library were processed similarly to those of the random library in (Ravarani et al., 2018). All processing steps aside from Illumina adapter sequence removal were completed using VSEARCH (Rognes et al., 2016). Forward and reverse read pairs were merged, allowing a maximum of one expected error when considering the quality scores per base. Adapter sequences were removed using cutadapt (Martin, 2011). Sequences were then deduplicated across all samples, counting the number of times each unique sequence appeared. The sequences were then filtered to include only those with a length of at least 60 bases and appearing at least twice across the library. The deduplicated sequences were clustered between those with a minimum sequence similarity of 90%, in an attempt to prevent two sequences with minor differences from being considered distinct sequences (Ravarani et al., 2018). These sequence clusters were then considered centroids, to which the original reads (merged, without adapters) were mapped and counted. For this step, the sequence identity parameter was set to 80%.

Reads from the design library were mapped using Kallisto(Bray et al., 2016), thanks to the improved alignment rate offered by the 20-nucleotide barcode present in each sequence of the library. Kallisto performs pseudoalignment, a probabilistic method, to map reads to the design library sequences and their barcodes, thereby providing the abundances of each sequence in the library. For each sample, cutadapt was used to remove adapters and demultiplex by the biorep barcode (Martin, 2011). Reads were then pseudoaligned to the design library (tAD sequence plus individual barcode) using Kallisto, with a default kmer size of 31.

### Estimation of sequence growth rates

Correlations among the read counts of biological replicates were calculated to ensure reasonable consistency, and sequences with at least five counts in at least two of the five biological replicates at baseline (identified by the five unique BioRep barcodes that were appended during PCR amplifications) were retained for subsequent analysis. Sequence counts were then averaged across biological replicates, resulting in one value for each sequence in each sample. These were then normalized within each sample (read counts divided by total reads in sample) to control for overall quantification differences between samples, and normalized to the baseline to quantify cell growth (by calculating the log2 fold change of each sequence at each time point versus its counts at time 0). The result of this step was a set of baseline-centered read counts for each sample at each of the five time points, which could then be plotted to determine whether abundance increases or decreases over time. Robust linear regression (implemented in the MASS package in R) was used to estimate the slope of each sequence over time; this was our final estimate for the functionality of each sequence (Venables et al., 2002). Regression of sequence counts vs. day was conducted from day 0 to 4 for most sequences, forcing a y intercept of 0 (because counts were normalized against day 0). Sequences for which the read count dropped and stayed below 3 were regressed from day 0 through the first day at which their read count was below 3. To define a strict binary cutoff for defining functional versus nonfunctional sequences, the 5 highest growth slopes out of 50 total stop codon sequences (the unique sequences that started with a stop codon) were averaged and used for individual sequence classification. For data visualization, all growth slopes were recentered to this cutoff slope so that the cutoff slope became zero.

### Sequence feature analysis

Sequence features such as the presence or number of individual amino acids or multiresidue motifs, the balance of aromatic versus acidic residues, and the mixing between amino acid residues were used in machine learning analyses. Balance was defined as the difference between the number of aromatic and acidic amino acids, while mixing was defined as the number of aromatic-acidic dipeptides plus the number of acidic-aromatic dipeptides. Ridge regression was conducted using the caret package in R (Kuhn, 2020). Ridge logistic regression was performed with a number of feature vectors for predicting functional vs non-functional ADs from amino acid sequences. Training and testing the ridge model was performed with the WD12 subset of the Gal4 design library (1130 functional and 2638 non-functional ADs). This subset was split into 5 even groups. Four were used for training and one was used as the testing set. The training/testing process was repeated five times using a different hold-out test set each time. ROC (receiver operating characteristic) curve AUC was computed on the test set after each training round, and averaged across the five rounds to find the final reported AUROC (Fig. 2G) for each of the following feature vectors. The aa composition: 2 features (# of W and # of D); aa x position: 24 features (2 amino acids [W,D] x 12 positions); tetrapeptides: 16 features (all tetrapeptides of W and D); tetrapeptide x position: 144 features (16 tetrapeptides x 9 positions); tetrapeptide x position + balance: 145 features (16 tetrapeptides x 9 positions) + (#W - #D).Neural network prediction of functionality was performed using the Keras and TensorFlow packages, where each sequence is transformed into a one-hot encoded 3 × 20 matrix (3 amino acids G, W, or D & 20 positions) (Chollet, 2015) (Abadi et al., 2015). The neural network architecture was simple, consisting of a flattened input matrix, two fully connected hidden layers of 60 and 30 nodes, with a dropout rate of 0.2 after each hidden layer, and finally connected to a softmax output layer predicting 1 – functional or 0 – nonfunctional.

### Structure prediction and analysis

Secondary structure prediction was performed with SPOT-1D (Singh et al., 2021). The predictions were calculated for each candidate 30-aa-long tAD sequence only. The SS3 output of SPOT-1D was then used to assign a helicity percentage to each candidate tAD sequence. The visualization of structure predictions utilized the AlphaFold2 Colab notebook (Jumper et al., 2021). Sequence structures were predicted for both the candidate tAD and the preceding “linker” (PEFVIRLTIGRAAIMEEQKLISEEDLHMAMG). Visualizations of the candidate tADs were finalized in PyMOL V 1.8; the common “linker” sequence was removed, and key amino acids were colored.

---

**Supplementary Figure 1.**
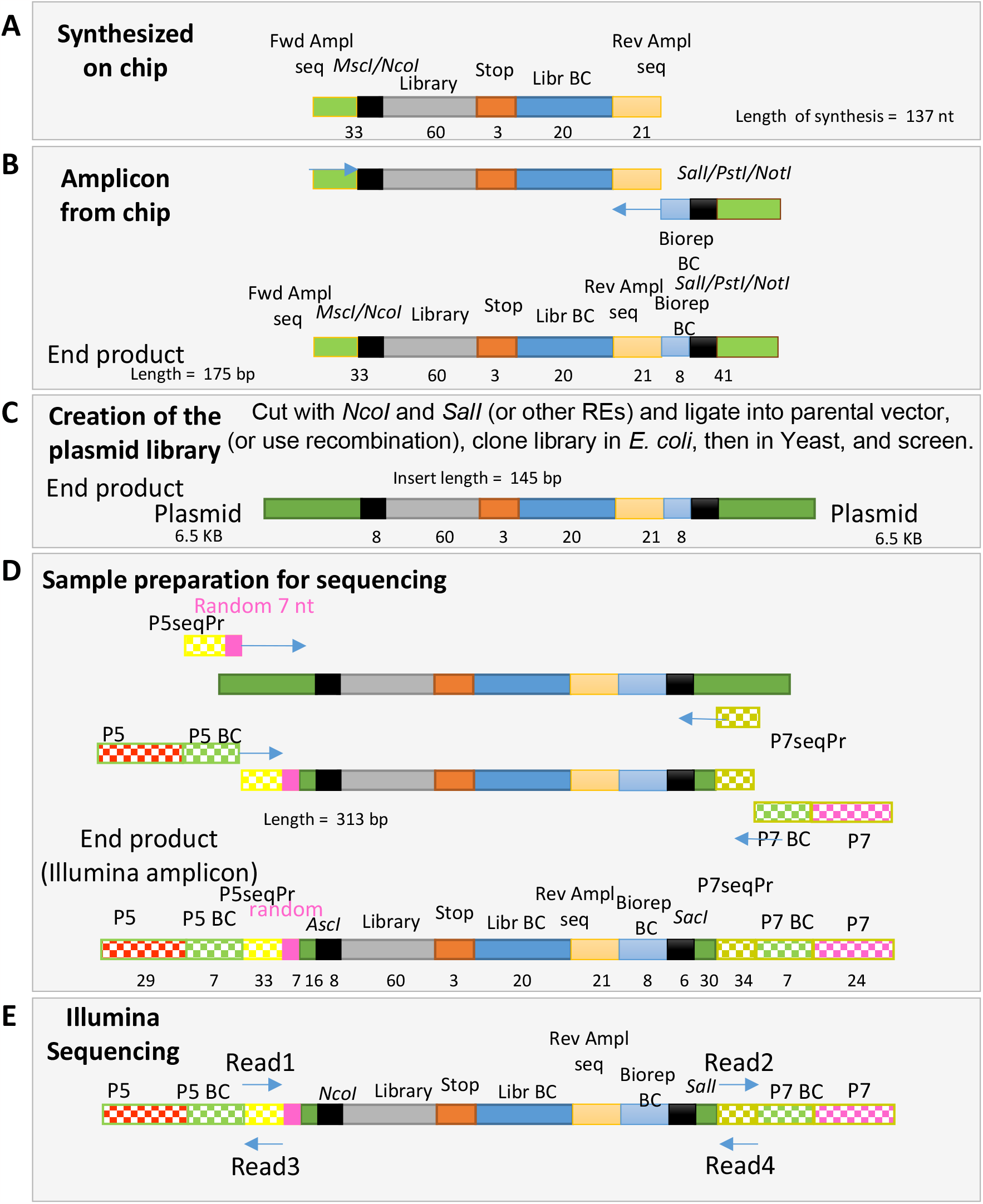
Schematic representation of wet lab steps: **A** – massively parallel synthesis of the design library; **B** – BioRep barcodes appending**; C** – cloning into parental yeast shuttle vector; **D** – sample preparation for NGS Illumina sequencing; **E** – sequencing at Illumina sequencing facility.

## Notes

### Competing Interest Statement

The authors have declared no competing interest.

